# Low temperature induced quiescence in parasitoids *Trichogramma evanescens* Westwood and *Trichogramma chilonis* Ishii reared on *Plodia interpunctella* (Hübner): Its utilization as quality control for mass rearing

**DOI:** 10.1101/2022.04.21.489033

**Authors:** Md. Mahbub Hasan, M. Nishat Parvin, Christos G. Athanassiou

## Abstract

Egg parasitoid, *Trichogramma* spp. is an important potential biological control agent for wide range of lepidopteran pests. Cold storage of host eggs has been proposed as a valuable technique for ensuring the release of sufficient parasitoid numbers whenever it is needed. Thus, the impact of low temperatures to induce quiescence in *Trichogramma evanescens* and *Trichogramma chilonis* was studied using eggs of *Plodia interpunctella* as hosts. Prepupae of the parasitoids *Trichogramma evanescens* and *T.chilonis* were stored for 15, 30, 45, 60 and 75 d at 4°C, following a 7 d period of acclimation at 10°C. Both parasitoid species seem to survive unfavorable temperature conditions by entering a state of quiescence. Parasitism, adult emergence, female proportion relative to male and F_1_ adult emergence were not affected by cold storage in either parasitoid for up to 30 d of storage. Parasitized host eggs of *P. interpunctella* can be stored for up to 60 d at 4°C for both parasitoids, but there was no emergence at 75 d. Results clearly show that there are specific intervals of cold storage during which the parasitoids can remain unaffected for relatively long periods of time. Although we observed some adverse effect in longevity and parasitism rates, the technique described here can be further utilized in mass rearing strategies of egg parasitoids for relatively long periods that will allow shipment and application in biological control programs.

## Introduction

Among the various insect egg parasitoids, *Trichogramma* (Hymenoptera: Trichogrammatidae) species are widely used for pest management throughout the world [1,2]. The most common species of this genus are *Trichogramma minutum* Riley, *T. evanescens* Westwood, *T. chilonis* Ishii, *T. pretiosum* Riley, *T. deion* Pinto and Oatman and *T. dendrolimi* Matsumura [3]. These species have been found in an extremely large variety of tritrophic systems, and on different hosts [4]. In general, *Trichogramma* spp. are among the most exploited groups of parasitoids for biological control and integrated pest management. So far, species of this genus have been considered in management programs for various insect pests on 18 crops and can either be explored as alternatives to chemical pesticides or as important components in integrated pest management (IPM) programs [5].

A period of dormancy interrupts the life cycle of many insect species, serving to protect them from harsh environmental conditions. Dormancy could occur in most of the parasitoids either by quiescence or diapause [6–9]. In quiescence, insect development is halted or slowed in direct response to unsuitable environmental conditions, while development resumes under favorable conditions, although this phenomenon varies among species [10,11]. In contrast with quiescence, diapause concerns the period of developmental arrest combined with low metabolic activity which is characterized by behavioural inactivity and slowing growth. In addition, diapause occurs at a species-specific stage of ontogenesis and its expression is regulated by various environmental signals [12,13]. The success of biological control programs is often related to problems and cost of rearing beneficial insects due to their relatively short shelf-life and difficulties in synchronizing parasitoid and host life cycles, and hence, dormancy can be used to mitigate this problem [14]. It has been reported that the dormancy management tools play an important role for biological control involving sterile insects or natural enemy releases [13]. The implementation of most suitable “parasitoid storage” methods is considered as a valuable tool to maintain high quality mass production and to allow synchronization of field releases during pest outbreaks [15,16]. In recent years, cold storage of host eggs received more attention since it can optimize the use of biological control agents [17]. Moreover, it provides the availability in maximum numbers at the time of release, along with flexibility and quality assurance in mass production [18]. In general, storage temperatures ranging from 0 to 15°Care widely used for parasitoids [15]; however, the optimal temperature depends upon the relative balance between the reduction of metabolic rates and the risk of accumulating chilling-related injury [16]. A gradual or rapid acclimation is usually desirable for pre-exposure to sublethal low temperatures [19–21]. Moreover, it has been reported that there is a positive impact on cold storage tolerance for acclimated parasitoids [22,23] despite the fact that there might be some effects in decreasing tolerance as well [24].

Cold storage of parasitoids is crucial for mass production, as the major outcome of this procedure is quality control of the exposed individuals [25–28]. In the case of species of the genus *Trichogramma*, storage can be achieved with or without previous diapause induction [29]. Smith [30] reported that immatures of several *Trichogramma* species can enter diapause or quiescence within host eggs and thereby tolerate long periods of subfreezing temperatures.

In this context, the objective of the present study was to determine the effect of storage of parasitized eggs at low temperature for varying periods on key biological traits that are directly related to parasitism and longevity in *T.evanescens* and *T. chilonis*, two egg parasitoids that have been thoroughly tested for the control of stored product species [31].

## Materials and methods

### Parasitoid and host rearing

The two parasitoids species were originally collected from vegetable field crops and maintained as stock culture at the Bangladesh Agricultural Research Institute (BARI), Gazipur and Sugarcane Research Institute, Ishwardi, Bangladesh. These species were cultured and mass-reared using eggs (<24h old) of the Indian meal moth, *Plodia interpunctella* (Hübner) (Lepidoptera: Pyralidae), at the Post-Harvest Entomology Laboratory, Department of Zoology, Rajshahi University, as suggested by Hegazi et al. [32]. The host eggs were pasted on paper strips (2 X 10 cm) with gum arabic glue (DalerRowney Co., India) and then exposed to the parasitoids in glass jars (2 L) supplied with cotton swabs soaked in honey solution(10%) for adult diet [33]. The jars were covered by cloth-wrapped cotton fixing with rubber bands. The egg strips were changed daily to minimize super-parasitism. The culture of Indian meal moth *P. interpunctella* was maintained at the Post-Harvest Entomology Laboratory, Department of Zoology, Rajshahi University, Bangladesh. They were reared on a standardized diet as suggested by Phillips and Strand [34]. The parasitoids and host cultures were maintained in an incubator set at 27°C, 70% relative humidity (RH), and a photoperiod of 16:8 (L: D) h.

### Experimental setup for cold storage

Fifteen pairs of *T. evanescens* and *T. chilonis* adults (1-2 d old) were kept separately in twenty glass tubes (84 X 2.5 cm) containing five host egg paper strips in each tube (2 X10 cm) for 24 h at 25°C, 16:8 (L:D) h and 60 to 70% RH [35](Doyon and Boivin, 2005). One hundred host eggs were pasted on each paper strip with gum arabic glue, and exposed to either of the species. The glass tubes were provided with cotton swabs soaked in 10% honey solution for adult diet. After exposure for 24 h, 50 paper strips containing parasitized eggs for each species were collected and transferred to acclimation conditions of 10°C, 8:16 L:D and 70% RH, for 7 d. After that, the parasitized egg paper strips were transferred to an incubator set at 4°C, 24 h dark and 60-70% RH for 0, 15, 30, 45, 60, and 75 d [28]. A control treatment with no acclimation/cold storage periods was maintained at standard rearing conditions of 25°C, 16:8 L:D and 70% RH [35]. Three replicates, having fifty parasitized eggs (blackened color) per strip for each parasitoid were carried out per treatment. After completing the desired storage periods, the egg strips were removed from each treatment and kept successively at 10, 15 and 20°C for 24h at each temperature. Finally, the egg strips were shifted to an incubator set at 25°C, 16:8 L: D and 70% RH until adult emergence. The experimental set up for cold storage is summarized in the schematic diagram of Fig. 1. After that, the percent emergence as well as sex-ratio (female proportion) for each parasitoid was recorded. In order to measure the longevity of emerging adults, the individuals were placed separately in tubes with diet (10% honey solution) [33] and were checked every day.

**Fig. 1.**
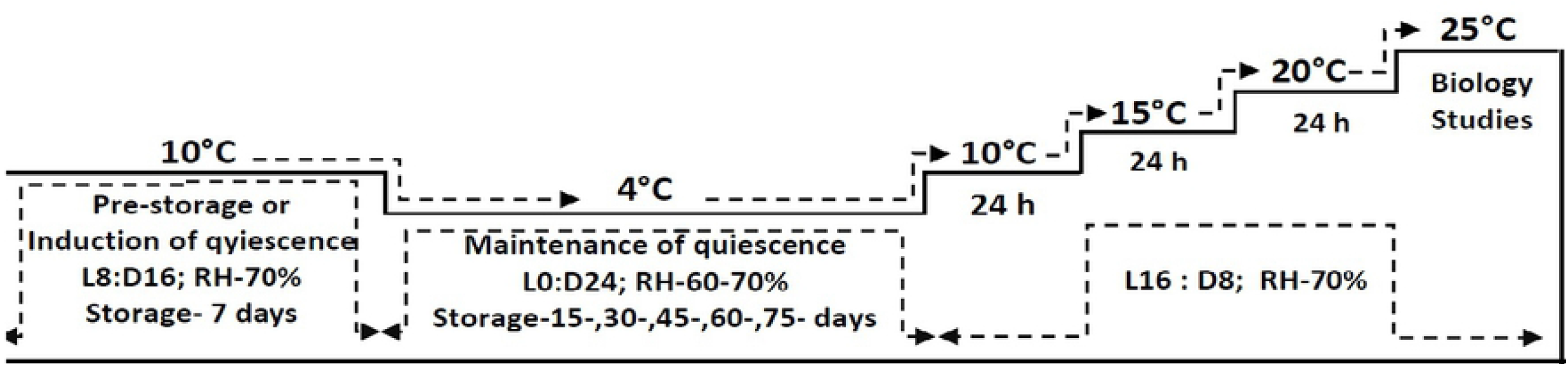
Schematic diagram of the experimental protocol for inducing quiescence in parasitoids.

In order to determine the effectiveness of parasitism, twenty five fresh host eggs (<24h old) were pasted on paper strips and kept in glass vial (84 X 2.5 cm) containing a pair of male and female adult parasitoids that have been developed from cold storage. There were three replicates having twenty five eggs per strip for each parasitoid and treatment. The egg strips were changed daily to minimize super-parasitism. The parasitism efficiency of parasitoid that have been developed from cold storage parasitized eggs was determined on the basis of general productivity (GP), reduction in parasitism efficiency (RPE), relative to control as RPE and parasitism efficiency (PE).The adult emergence percentage was calculated as number of emerged adults vs. the number of parasitized eggs x 100. The general productivity (GP) was estimated as GP = rate of emergence x rate of produced females in progeny x number of parasitized eggs [36], and reduction in parasitism efficiency (RPE) was determined in relative to control as RPE = [(GP of control batch–GP of individual storage exposure)/ GP of control batch] x 100 while parasitism efficiency (PE) was calculated as PE = (GP of individual storage exposure/GP of control batch) x 100.

### Statistical analysis

The validation of the data distribution of each measured variables was carried out using Shapiro–Wilk’s test. The data for percent values were arcsine transformed before the analysis (untransformed data are presented in the Figures and Tables). Then, all data were analyzed through a two-way factorial analysis of variance (ANOVA) using the PROC ANOVA [37], separately, for each of the trait including parasitism rates, percent emergence, female proportion and adult longevity. Means were compared by Tukey-Kramer HSD test at 5%.

## Results

### Experimental setup for cold storage

The durations of cold storage of parasitized eggs of *P. interpunctella* significantly affected the percent adult emergence in both *T. evanescens* (F=45.36; df=5,12; P<0.001) and *T. chilonis* (F=44.18; df=5,12; P<0.001). Moreover, there was also a significant difference between species in adult emergence (F=28.67; df=1,24; P<0.001). In the control, the maximum percent of parasitoid emergence reached 76 and 81% for *T. evanescens* and *T. chilonis*, respectively, while the minimum was 28% for the 45d storage at 4°C for *T. evanescens* (Fig. 2). However, no adult emergence was recorded from the 75 d cold storage periods for either species. In general, the overall percent emergence was significantly higher (F=54.85; df=6,29; P<0.001) in *T. chilonis* compared to *T. evanescens* in all the cold storage periods with the exception of 15 d (Fig. 2). There was also no significant (F=3.30; df=5,24; P=0.23) difference between the interaction of species*storage in percent adult emergence. The proportion of female relative to male (F/M ratio) resulting from the cold-stored host eggs did not vary significantly in *T. evanescens* (F=2.64; df=6,8; P=0.10) and *T. chilonis* (F=2.63; df=6,8; P=0.10) (Fig. 3). The proportion of female also did not differ significantly between species (F=1.28; df=1,5; P=0.27). Longevity of females significantly declined with increase of exposure of cold storage in both *T. evanescens* (F=56.03; df=5,12; P<0.001) and *T. chilonis* (F=56.38; df=5,12; P<0.001) (Fig. 4). The adult longevity did not differ significantly between the species (F=2.25; df=1,24; P=0.15) produced from cold stored host eggs and it implies the same trend in the interaction of specie*storage (F=1.50; df=5,24; P=0.23).

**Figure 2:**
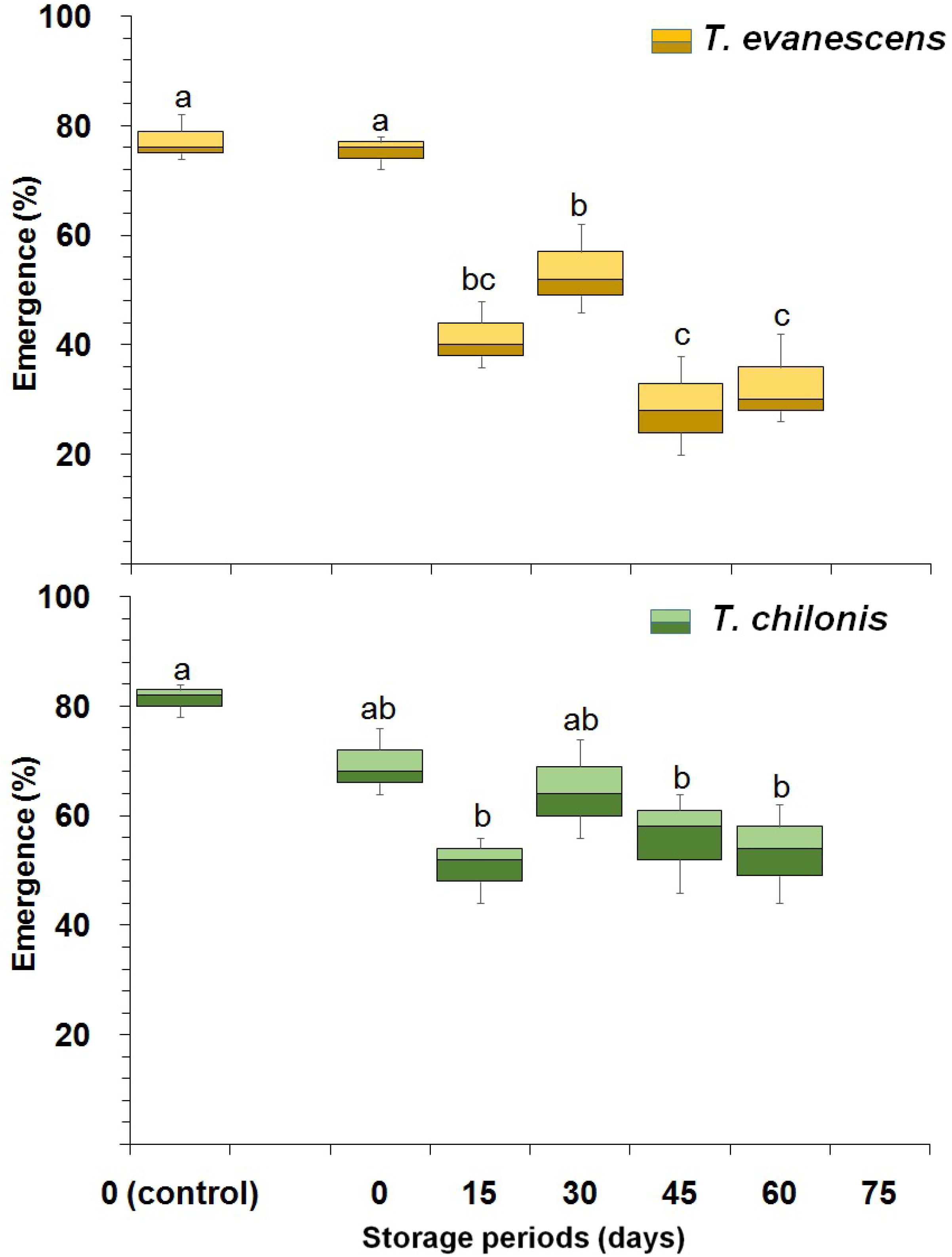
Box plot for the effect of cold storage period on the percentage of adult emergence of *T. evanescens* and *T. chilonis* reared on *P. interpunctella*. [Boxes depict median and quartiles (75 and 25% scores), whiskers depict upper and lower limits; Bars within each species followed by the same letters are not significantly different; HSD test at 0.05; Lowercase for *T. evanescens* and uppercase for *T. chilonis*]

**Figure 3:**
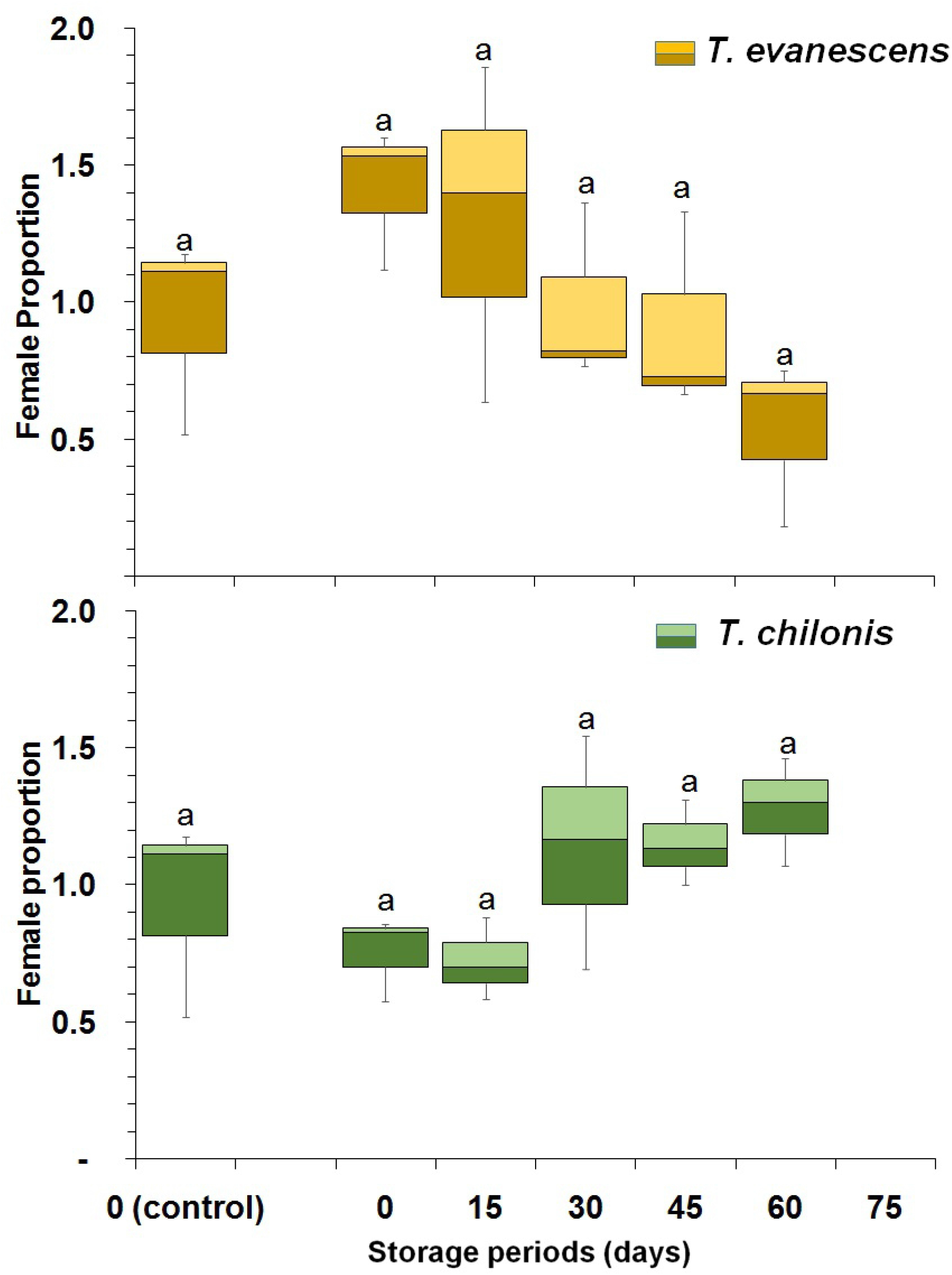
Box plot for the effect of cold storage period on the female proportion of *T. evanescens* and *T. chilonis* reared on *P. interpunctella*. [Boxes depict median and quartiles (75 and 25% scores), whiskers depict upper and lower limits; Bars within each species followed by the same letters are not significantly different; HSD test at 0.05; Lowercase for *T. evanescens* and uppercase for *T. chilonis*]

**Figure 4:**
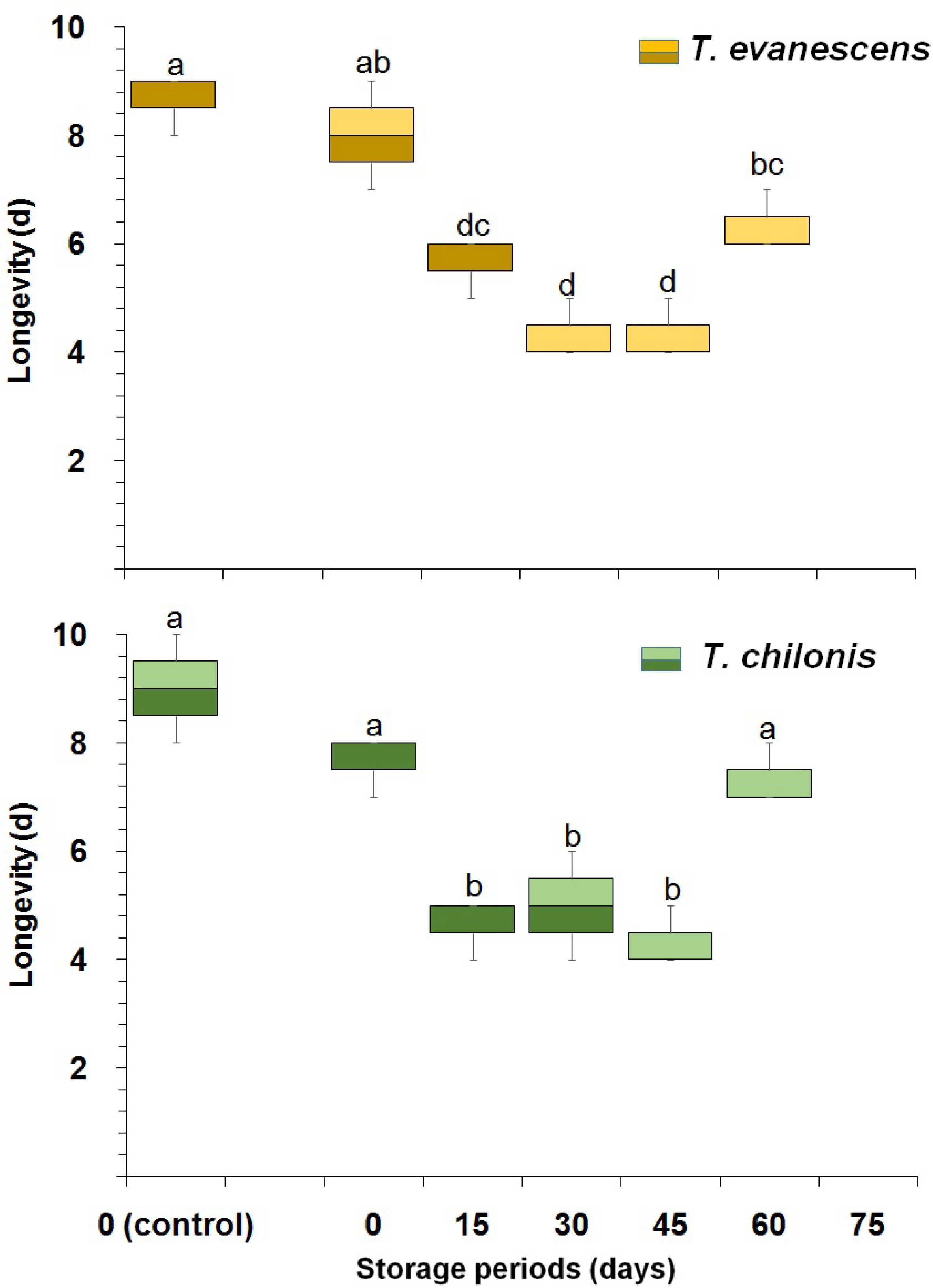
Box plot for the effect of cold storage period on the adult longevity of *T. evanescens* and *T. chilonis* reared on *P. interpunctella*. [Boxes depict median and quartiles (75 and 25% scores), whiskers depict upper and lower limits; Bars within each species followed by the same letters are not significantly different; HSD test at 0.05; Lowercase for *T. evanescens* and uppercase for *T. chilonis*]

Parasitism percent of adult females in F_1_ progeny were higher in non-cold stored host eggs (control) for *T. evanescens* (81%) and *T. chilonis* (86%), as compared with the cold stored ones (Fig. 5). Results indicated that cold storage of host eggs had a significant effect on the F_1_ parasitism of *T. evanescens* (F=21.17; df=5,12; P<0.001) and *T. chilonis* (F=22.51; df=5,12; P<0.001) females in the F_1_ generation. There were also significant differences (F=4.33; df=1,24; P<0.04) between species in F_1_ parasitism. The percentage of parasitism decreased in adult females of *T. evanescens* and *T. chilonis* resulting from cold stored host eggs compared to non-cold stored ones. Moreover, females emerging from 0 and 30 d cold stored host eggs showed better parasitism performance compared to other cold storage durations (Fig. 5). The percentage of adults emerged in the F_1_ progeny was significantly affected by the cold storage in both the species *T. evanescens* (F=7.13; df=5,12; P<0.002) and *T. chilonis* (F=4.53; df=5,12; P<0.01) (Fig 6). The highest percentage (83%) of adult emergence in F_1_ was noted for *T. evanescens* that developed from 15 d cold storage duration compared to others and there was also significant (F=6.61; df=1,29; P<0.02) variation between the species (Fig 6).

**Figure 5:**
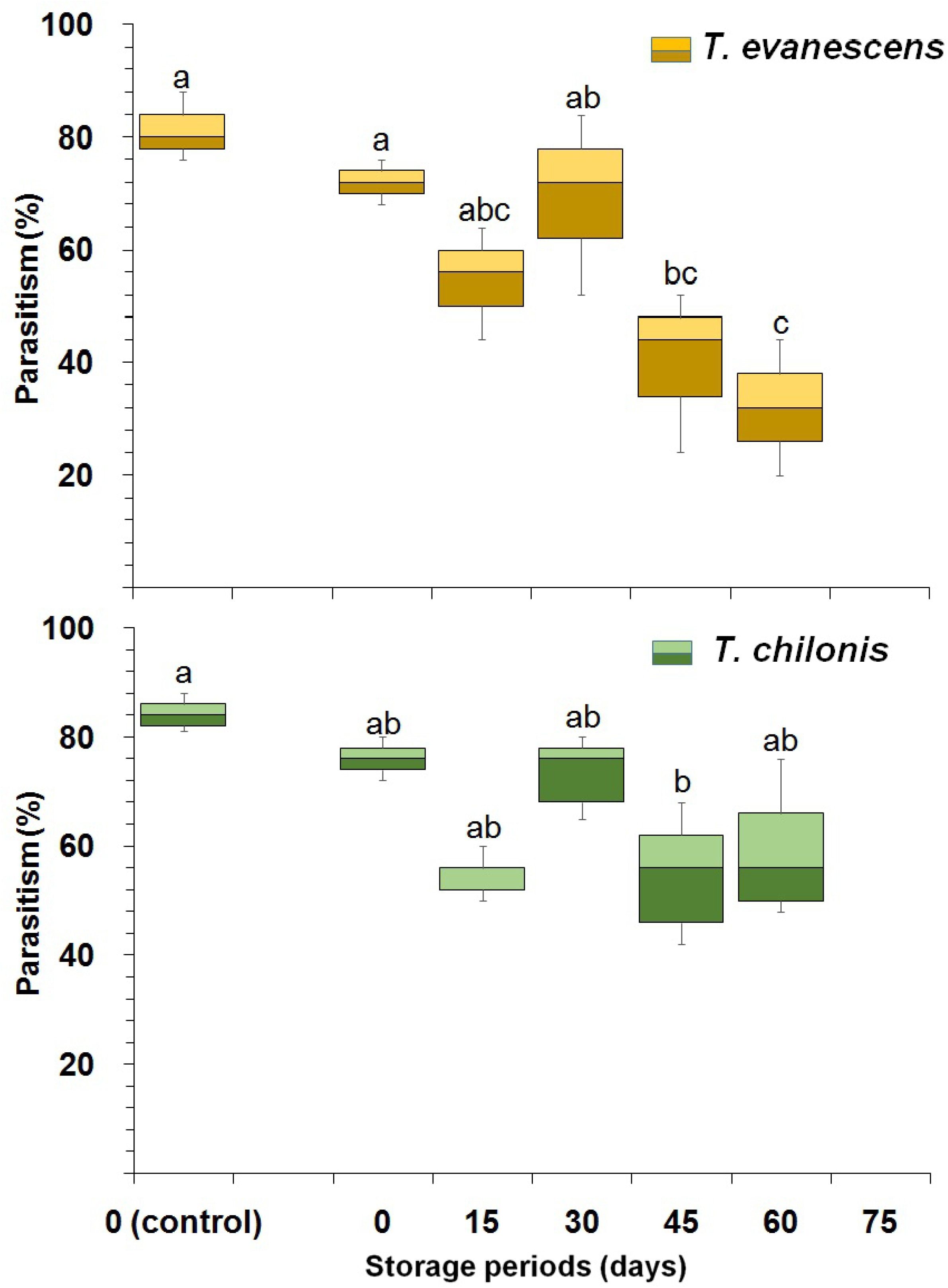
Box plot for the effect of cold storage period on the percent parasitism in *T. evanescens* and *T. chilonis* reared on *P. interpunctella*. [Boxes depict median and quartiles (75 and 25% scores), whiskers depict upper and lower limits; Bars within each species followed by the same letters are not significantly different; HSD test at 0.05; Lowercase for *T. evanescens* and uppercase for *T. chilonis*]

**Figure 6:**
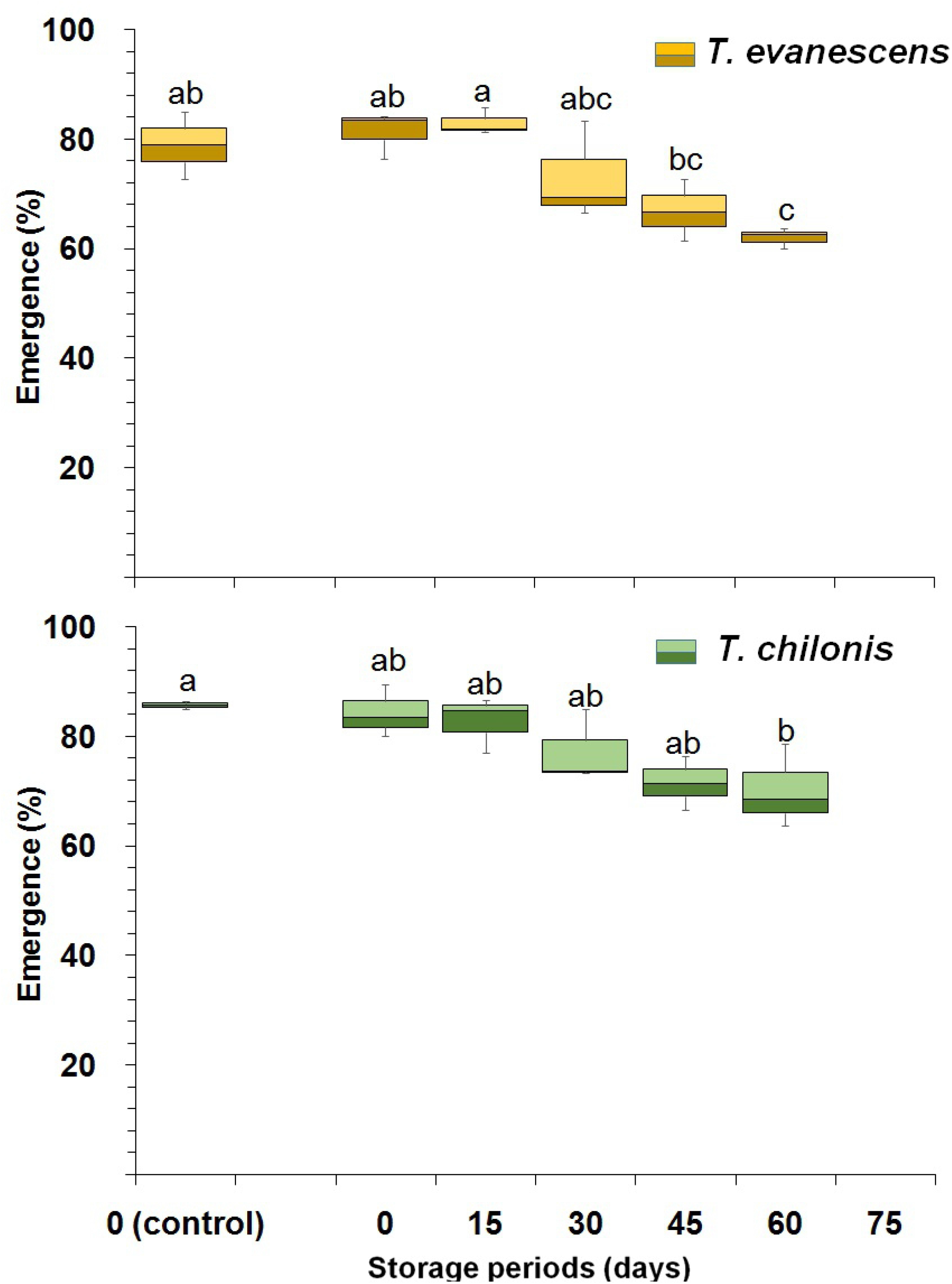
Box plot for the effect of cold storage period on the percent F_1_ adult emergence in *T. evanescens* and *T. chilonis* reared on *P. interpunctella*. [Boxes depict median and quartiles (75 and 25% scores), whiskers depict upper and lower limits; Bars within each species followed by the same letters are not significantly different; HSD test at 0.05; Lowercase for *T. evanescens* and uppercase for *T. chilonis*]

In general, key biological traits, such as fecundity (blackened eggs), adult emergence and ratio of female, of the parasitoid adults that emerged from host eggs that had been in cold storage were reduced as compared to controls (Table 1). The values of general productivity (GP) gradually decreased significantly as duration of storage lengthened for both species (*T. evanescens:* F=9.22; df=5,12; P<0.001 and *T. chilonis:* F=12.77; df=5,12; P<0.001) (Table 1). However, parasitism efficacy (PE) was significantly reduced up to 30.96 and 26.95 for *T. evanescens* (F=7.94; df=4,10; P<0.003) and *T. chilonis* (F=6.31; df=4,10; P<0.008) respectively at the 60 d cold storage interval. Accordingly, the highest reduction in parasitism efficiency (RPE) reached up to 69.04 and 73.05 %, at 60 d cold storage groups for *T. evanescens* and *T. chilonis* which also varied significantly (*T. evanescens*, F=7.94; df=34,10; P<0.003 and *T. chilonis*, F=6.31; df=4,10; P<0.008) respectively (Table 1).

**Table 1:**
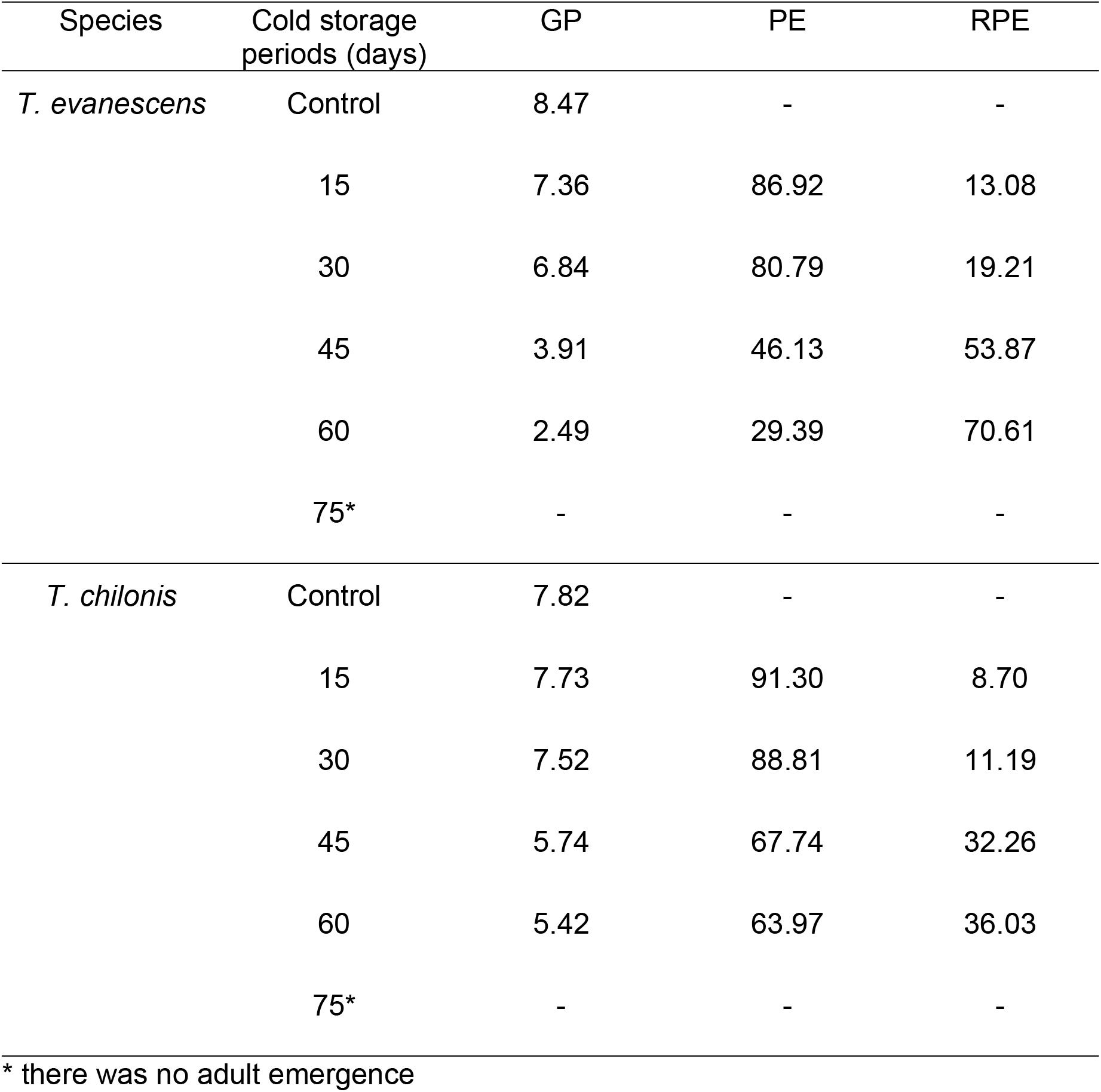
General productivity (GP), parasitism efficiency (PE), and reduction in parasitism efficiency (RPE) of *T.evanescens* and *T. chilonis* reared on parasitized *P. interpunctella* eggs that had been stored at 4°C for different periods

| Species | Cold storage periods (days) | GP | PE | RPE |
| --- | --- | --- | --- | --- |
| *T. evanescens* | Control | 8.47 | - | - |
|  | 15 | 7.36 | 86.92 | 13.08 |
|  | 30 | 6.84 | 80.79 | 19.21 |
|  | 45 | 3.91 | 46.13 | 53.87 |
|  | 60 | 2.49 | 29.39 | 70.61 |
|  | 75* | - | - | - |
| *T. chilonis* | Control | 7.82 | - | - |
|  | 15 | 7.73 | 91.30 | 8.70 |
|  | 30 | 7.52 | 88.81 | 11.19 |
|  | 45 | 5.74 | 67.74 | 32.26 |
|  | 60 | 5.42 | 63.97 | 36.03 |
|  | 75* | - | - | - |

\* there was no adult emergence

## Discussion

The results of the current study clearly suggest that cold storage of *P. interpunctella* eggs for 30d is a viable storage duration for production of increased quantities of *Trichogramma* parasitoids. At the same time, this storage maintains also the “quality” of the parasitoids, i.e. their ability to provide satisfactory progeny production and the concomitant increased parasitism rates. Our study shows that storing *P. interpunctella* eggs at 4 °C more than 60d, negatively influences the overall survival, and should be avoided.

It is not clear if the exposed individuals here entered diapauses or the experimental protocol followed here caused quiescence, given that quantitative data were not collected towards this direction. This direct response to non-suitable environmental conditions, and the subsequent response with the return of the favorable conditions, i.e. increase of temperature and host availability, may mostly suggest quiescence rather than diapause [8,38,39]. Andreadis et al. [39] found that the parasitoid was generally more susceptible than the host while following a similar protocol with low temperature or the exposure of the endoparasitoid *Venturia canescens* (Gravenhorst) (Hymenoptera: Ichneumonidae) on larvae of the Mediterranean flour moth, *Ephestia kuehniella* Zeller (Lepipdoptera: Pyralidae). However, in that study, the age of the parasitoid played a critical role in its survival after exposure to low temperature, which can be taken into account to reduce mortality in cases of low temperature parasitoid preservation [35]. In our case, we did not measure survival of the host, in terms of *P. interpunctella* egg hatch, but our data show that both *T. evanescens* and *T. chilonis* were able to survive long periods at low temperature, at percentages of emergence that were generally lower than those of individuals that were not exposed to cold, suggesting that even an exposure as short as 15 d can cause some parasitoid morality. It is generally considered that exposure, and thus preservation from a practical point of view, of parasitoids at low temperature may be subjected to considerable fitness costs, including increased mortality [16,38,39], which stands in accordance with the present findings. However, Cagnotti et al. [38] reported that *Trichogramma nerudai* Pintureau (Hymenoptera: Trichogrammatidae) prepupae, that had been stored at suboptimal temperature were not affected when the storage was 10 or 20d. In this context, preconditioning may play a positive role [38,39].

Despite some mortality after exposure to cold, we found that these losses are partially compensated by specific adaptations that were not expected, such as changes to male: female ratios, for both species, and progeny production attributes that are similar to those in the case of the un-acclimated individuals. This is expressed more vigorously in the case of *T. chilonis*, which generally performed better than *T. evanescens*. Interestingly, despite the fact that exposure to cold reduced longevity of both species for exposure periods of 15, 30 and 45 d, at 60 d longevity was increased to levels that were comparable to controls. This may be attributed to the initiation of a cold-hardening procedure that might have been triggered at these conditions, switching eventually the adaptation of the parasitoids. The increased adaptation, through improved cold-hardening that is exposed more vigorously in long exposures, has been also identified in other parasitoids [38,40].

For the endoparasitoid *Microplitis prodeniae* Rao and Chandry (Hymenoptera: Braconidae), Yan et al. [41] found that there were no subsequent effects of exposure to cold of the parental individuals during the two next generations. Similar results have been also reported by Silva et al. [42] for the aphid parasitoid *Diaeretiella rapae* (McIntosh) (Hymenoptera: Braconidae). It is well established that exposure to cold is likely to cause some major, and in most cases irreversible, negative changes in fecundity of the exposed parasitoids, which, in turn, may seriously affect their performance after termination of the exposure [42–44]. For instance, Colinet and Hance [45] found that exposure to suboptimal temperatures seriously affected the reproductive system of the aphid parasitoid *Aphidius colemani* (Viereck) (Hymenoptera: Braconidae). In accordance with what has been reported by Yan et al. [41] for *M. prodeniae*, parasitism rates of the offspring that were produced by the exposed parental adults were not seriously affected, suggesting that any effect is physiologically-related and not transferable to the next generation. This is particularly important in developing strategies for cold storage of parasitoids, and most studies suggest that there is no carryover to the subsequent generations that will be produced by the exposed parental individuals. Still, our work is in accordance with previous reports considering the negative effect of exposure to low temperature to progeny production capacity, demonstrated in both *T. evanescens* and *T. chilonis*. Paradoxically, we found that the relationship between parasitism rates and duration of exposure to cold was not linear, and, to some extent, gave dissimilar results. While for *T. evanescens* exposure for 15 and 30 d gave significantly similar parasitism results with the controls, longer exposures was subsequently reduced to levels that were not comparable to those of the controls. On the other hand, for *T. chilonis*, parasitism caused by parasitoid adults that had been exposed for 15 d was lower than that in the controls, but at longer exposures did not differ significantly from controls. All the above suggest the potential occurrence of specific physiological responses that are exhibited only at specific temperature-exposure combinations, and merit additional investigation. However, our results can be also related to the effect of low temperature on *P. Interpunctella* eggs, a feature that might have indirectly affected survival and performance of the parasitoids. In this context, Athanassiou et al. [46] found that eggs of *P. interpuntella* were particularly susceptible to cold, and were much more susceptible than larvae, in contrast with most stored product insects where the egg is by far the most cold tolerant life stage [34,46,47]. The exposure of eggs to cold may cause temporal physical damage to host eggs, that subsequently influences the overall parasitoid performance.

## Conclusion

The results of our work clearly show that there are specific intervals of cold storage during which the parasitoids can remain unaffected for relatively long periods of time, which here can reach 60 d, if the storage temperature is +4°C. Although we observed some adverse effect in longevity and parasitism rates, the technique described here can be further utilized in mass rearing strategies of egg parasitoids for relatively long periods that will allow shipment and application in biological control programs. The cost of cold storage of parasitoids should be further analyzed, in comparison with additional storage methods that have been proposed.

## CRediT authorship contribution statement

**Md. Mahbub Hasan:** Conceptualization, Methodology, Writing - original draft, Funding acquisition. **M. Nishat Parvin:** Methodology, Investigation, **Christos G. Athanassiou:** Conceptualization, Writing – final draft, review & editing.

## Acknowledgements

We are grateful to David Denlinger (Department of Entomology, Ohio State University) for comments and suggestion on an earlier version of this paper. We thank to the Bangladesh Agriculture Research Institute (BARI), Gazipur and Sugarcane Research Institute, Ishwardi, Bangladesh for providing the parasitoids. The authors are grateful to the Department of Zoology, Rajshahi University for extending the laboratory facilities.

## Authors’ contributions

The authors cooperated in all the experiments, statistical analysis of data, reading, and approval of the final manuscript.

## Ethics approval and consent to participate

Not applicable.

## References

1. Cagnotti CL, Lois M, Silvia NLP, Botto EN, Viscarret MM. Cold storage of *Trichogramma nerudai* using an acclimation period. BioControl. 2018; 63: 565–573.

2. Siam A, Zohdy NZM, ELHafez AMA, Moursy LE, Sherif HAEL. Effect of different cold storage periods of rearing host eggs on the performance of the parasitoid *Trichogramma evanescens* (Westwood) (Hymenoptera: Trichogrammatidae). Egypt J Biol Pest Control 2019; 29:1–4.

3. Sumer F, Tuncbilek AS, Oztemiz S, Pintureau B, Rugman-Jones P, Stouthamer R. A molecular key to the common species of *Trichogramma* of the Mediterranean region. BioControl 2009; 54: 617–624.

4. Consuelo MDeM, Lewis WJ, Tumlinson JH. Examining plant-parasitoid interactions in tritrophic systems. An Soc Entomol Bras. 2000; 29: 189–203.

5. Kumar P, Sekhar JC, Kaur J. Trichogrammatids: Integration with Other Methods of Pest Control. In: Biological Control of Insect Pests Using Egg Parasitoids, (Eds.) Sithanantham S, Ballal CR, Jalali SK, Bakthavatsalam N, Springer Paris, 2013;191–208.

6. Denlinger DL. Dormancy in tropical insects. Ann Rev Entomol. 1986; 31: 239–364.

7. Rossi MM. Etude bioécologique des parasitoïphages *Trichogramma cacoeciae* Marchalet *T. evanescens* West. (Hym.,Trichogrammatidae) et du parasitoïdenymphal *Dibrachys affinis* Masi (Hym., Pteromalidae) associe’s a’ *Lobesia botrana* Den. And Schiff. (Lepidoptera: Tortricidae). PhD Thesis, Université de Rennes I, U.F.R. Sciences et Philosophie. 1993.

8. Boivin G. Overwintering Strategies of egg parasitoids, In: Biological Control with Egg Parasitoids (CABI, Wallingford), 1994; 219–244.

9. Denlinger DL. Regulation of diapause. Ann. Rev. Entomol. 2002; 47: 93–122.

10. Saunders DS. Insect clocks, 2nd edn. 1982; Pergamon Press, Oxford.

11. Chapman RF. The insects: structure and function, 4^th^ edn. 1998; Cambridge University Press, Cambridge.

12. Tougeron K. Diapause research in insects: historical review and recent work perspectives. Ent Exp et Applic. 2019; 167: 27–36.

13. Hasan MM, Hasan MM, Khatun R, Hossain MA, Athanassiou CG, Bari MA.. Mating attributes relating to parasitization and productivity in *Habrobracon hebetor* (Hymenoptera: Braconidae) rearing on host Indian meal moth (Lepidoptera: Pyralidae). J Econ Entomol. 2020; 113: 1528–1534.

14. Glenister CS, Hoffmann MP. Mass-reared natural enemies: scientific, technological, and informational needs and considerations. In: Proceedings Mass-reared natural enemies: application, regulation, and needs. (Eds) Ridgway, R., M.P. Hoffmann, M.N. Inscoe, and C.S. Glenister, Thomas Say Publications in Entomology, Entomological Society of America, Lanham, 1998; 242–247

15. Van Lenteren J, Tommasini M. Mass production, storage, shipment and quality control of natural enemies. In: Mass production, storage, shipment and quality control of natural enemies, integrated pest and disease management in greenhouse crops. (Eds.) Albajes, R., M.L. Gullino, J.C. van Lenteren and Y. Elad, 2002; 276–294. Springer, Dordrecht, Germany.

16. Colinet H, Boivin G. Insect parasitoids cold storage: a comprehensive review of factors of variability and consequences. Biol Control 201;158: 83–95.

17. Huang JHQ, Hua L, Wang Y, Zhang F, Li YX. Number of attacks by *Trichogramma dendrolimi* (Hymenoptera: Trichogrammatidae) affects the successful parasitism of *Ostrinia furnacalis* (Lepidoptera: Crambidae) eggs. Bull Ent Res. 2017; 107: 812–819.

18. Ghosh E, Ballal CR. Diapause induction and termination in Indian strains of *Trichogramma chilonis* (Hymenoptera: Trichogrammatidae). Can Entomol. 2017; 149: 607–615.

19. Chown SL, Nicolson SW. Insect physiological ecology: mechanisms and patterns. Oxford University Press, Oxford. 2004.

20. Chown SL, Terblanche JS Physiological diversity in insects: ecological and evolutionary contexts. Adv Insect Physiol. 2006; 33: 50–152.

21. Anguilletta MJ Jr. Thermal adaptation: a theoretical and empirical synthesis. Oxford University Press, Oxford, 2009.

22. Marwan IA, Tawfiq MM. Response of *Aphidius matricariae* Haliday (Hymenoptera: Aphidiidae) from mummified *Myzus persicae* Sulzer (Homoptera: Aphididae) to short term cold storage. Intl Pest Control 2006; 48: 262–265.

23. Luczynski A, Nyrop JP, Shi A. Influence of cold storage on pupal development and mortality during storage and on post-storage performance of *Encarsia formosa* and *Eretmocerus eremicus* (Hymenoptera: Aphelinidae). Biol Control 2007; 40:107–117.

24. Bernardo U, Iodice L, Sasso R, Pedata PA. Effects of cold storage on *Thripobius javae* (= *T. semiluteus*) (Hymenoptera: Eulophidae). Biocontrol Sci Technol 2008; 18: 921–933.

25. Hany AS, Abd El-Gawad ATEF, Sayed MM, Ahmed SA. Impact of cold storage temperature and period on performance of *Trichogramma evanescens* Westwood (Hymenoptera: Trichogrammatidae). Australian J Basic Appl Sc. 1991; 4: 2188–2195.

26. Leopold RA. Cold storage of insects for integrated pest management. In: Temperature sensitivity in insects and application in integrated pest management. (Eds) Hallman, G.J., and D.I. Denlinger, 1998; pp. 235–267, West View Press, Boulder, CO, USA.

27. Özder N. Effect of different cold storage periods on parasitization performance of *Trichogramma cacoeciae* (Hymenoptera, Trichogrammatidae) on eggs of *Ephestia kuehniella* (Lepidoptera, Pyralidae). Biocontrol Sci Tech. 2004; 14: 441–447.

28. Filho SRP, Leite GLD, Soares MA, Alvarenga AC, Paulo PD, Santos LDT, Zanuncio JC. Effects of duration of cold storage of host eggs on percent parasitism and adult emergence of ten Trichogrammatidae species. Fla Entomol 2014; 97:14–21.

29. Greenberg SM, Nordund DA, King EG. Mass production of *Trichogramma* spp.: Experience in the former Soviet union, China, United States and Western Europe. Biocontrol News Inform. 1996; 17(3): 51–60.

30. Smith SM. Biological control with *Trichogramma*: advances, successes, and potential of their use. Ann. Rev. Entomol. 1996; 41: 375–406.

31. Athanassiou CG, Arthur FH. Bacterial Insecticides and Inert Materials. In: Recent Advances in Stored Product Protection (Eds. Athanassiou, C.G., and F.H. Arthur). 2018; pp. 83, Springer-Verlag GmbH Germany.

32. Hegazi E, Adler C, Khafagi W, Agamy E.. Host-preference and parasitic capacity of new candidates of *Trichogramma* species (Hym.:Trichogrammatidae) against some stored product moths. J. stored Prod. Res. 2019; 80: 71–78.

33. Abd EL Hafez, A.. A comparison of thermal requirements and some biological aspects of *Trichogramma evanescens* (Westwood) and *Trichogrammatoidea bactrae* (Najgaja) reared from eggs of the pink and spiny bollworm. Ann. Agric. Sci. Ain. Shams. Univ. Cairo. 1995; 4: 901–912.

34. Phillips TW, Strand MR. Larval secretions and food odors affect orientation in female *Plodia interpunctella*. Entomol. Exp. Appl. 1994; 71: 185–192.

35. Doyon J, Boivin G. The effect of development time on the fitness of female *Trichogramma evanescens*. J. Insect Sci. 2005; 5:4, insectscience.org/5.4

36. Tshernyshev WB, Afonina VM. Optimal light and temperature conditions for *Trichogramma evanescens* Westwood rearing. In: Trichogramma and other Egg Parasitoids, 1995; pp 173–175, Fourth International Symposium Cairo, Egypt, 4–7 October 1994. INRA Publ, Paris.

37. SAS Institute. SAS/STAT 9.2 User’s guide. SAS Institute, Cary,North Carolina, USA. 2008.

38. Cagnotti CL, Lois M, Silvia NLP, Botto EN, Viscarret MM. Cold storage of *Trichogramma nerudai* using an acclimation period. BioControl 2018; 63: 565–573.

39. Andreadi SS, Spanoudis CG, Athanassiou CG, Savopoulou-Soultani M. Factors influencing super cooling capacity of the koinobiont endoparasitoid *Venturia canescens* (Hymenoptera: Ichneumonidae). Pest Manag Sci. 2013; 70: 814–818.

40. Reznik SY. Ecological and evolutionary aspects of photothermal regulation of diapause in Trichogrammatidae. J. Evol. Biochem. Phys. 2011; 47: 512–523.

41. Yan Z, Yue JJ, Bai C, Peng ZQ, Zhang CH. Effects of cold storage on the biological characteristics of *Microplitis prodeniae* (Hymenoptera: Braconidae). Bull Ent Res. 2017; 107: 506–512.

42. Silva RJ, Cividanes FJ, Pedroso EC, Barbosa JC, Matta DH, Correia ET, Otuka AK. Effect of low-temperature storage on *Diaeretiella rapae* (McIntosh) (Hymenoptera: Braconidae). Neotrop Entomol 2013; 42: 527–533.

43. Lacoume S, Bressac C, Chevrier C. Sperm production and mating potential of males after a cold shock on pupae of the parasitoid wasp Dinarmus basalis (Hymenoptera: Pteromalidae). J Insect Physiol. 2007; 53: 1008–1015.

44. Hance T, Baaren van J, Vernon P, Boivin G. Impact of extreme temperatures on parasitoids in a climate change perspective. Ann Rev Entomol. 2007; 52: 107–126.

45. Colinet H, Hance T. Male reproductive potential of *Aphidius colemani* (Hymenoptera: Aphidiinae) exposed to constant or fluctuating thermal regimens. Environ Entomol. 2009; 38: 242–249.

46. Athanassiou CG, Phillips TW, Wakil W. Biology and control of the khapra beetle, *Trogoderma granarium*, a major quarantine threat to global food security. Ann Rev Entomol. 2019; 64: 131–148.

47. Fields PG. Control of insects in post-harvest: low temperature. In: Physical Control Methods in Plant Protection, (Eds) Vincent, C., B. Panneton and F. Fleurat-Lessard, Springer, Paris, 2001; pp. 95–107.

